# Ribosomal proteins are major substrates of starvation-induced endosomal microautophagy in *Drosophila*

**DOI:** 10.64898/2026.09.01.748611

**Authors:** Prasoon Jaya, Satya Surabhi, Jennifer Aguilan, Simone Sidoli, Andreas Jenny

## Abstract

Maintenance of cellular homeostasis requires tight coordination between protein synthesis and degradation, particularly at old age and under conditions of stress including starvation. Autophagy contributes to sustain this balance by degrading cytoplasmic proteins and organelles. It thus is essential to prevent the accumulation of damaged proteins and organelles and to recycle nutrients. Of the three forms of autophagy, macroautophagy, chaperone mediated autophagy, and (endosomal) microautophagy (e-MI), the latter remains the least well understood. During e-MI, cytosolic substrate proteins are captured into late endosomes via ESCRT-dependent multivesicular body formation and then degraded in late endosomes or lysosomes. e-MI is thought to contribute to protein quality control under basal conditions and under stress. Importantly, very little is known about the endogenous substrates of e-MI in flies and thus about its physiological role. Performing integrative multi-omic analyses in *Drosophila* larval fat body that has functions similar to mammalian liver and adipose tissue, we identified 153 high-confidence endogenous e-MI substrates with the degradation of ribosomal proteins by e-MI being the most strongly affected functional category. Generally, we found that starvation caused the depletion of proteins involved in translation, aminoacyl-tRNA synthesis, and ribosomal biogenesis, without affecting their level of transcripts. Importantly, we observe a striking specificity between e-MI and macroautophagy, as the two pathways largely target distinct protein sets including different subsets of ribosomal proteins. Our metabolomic analysis further shows that genetic inhibition of e-MI reverses the reduced levels of amino acid caused by starvation. Together, our findings reveal ribosome turnover as a central physiological function of *Drosophila* e-MI and establish e-MI as a pathway driving metabolic adaptation during starvation.

**HIGHLIGHTS:**

- Proteomic and metabolomic analysis of *Drosophila* fat body under starvation stress
- Systematic identification of endosomal microautophagy (e-MI) substrates.
- Ribosomal proteins are a major target of e-MI.
- e-MI and macroautophagy degrade distinct proteins incl. different ribosomal ones.
- Prolonged starvation alters amino acid levels.

## INTRODUCTION

Cells maintain proteome homeostasis through the careful balance of protein synthesis and degradation, both of which are dynamically regulated to meet the constantly changing demands of an organism. Dysregulation of protein homeostasis leads to various human pathologies, including infectious diseases, cancer, and neurodegenerative disorders that increase with aging (1–3). Autophagy is the lysosomal degradation of cytoplasmic components. While short-lived proteins are mostly degraded by the proteasome, autophagy is responsible to degrade and recycle long-lived, dysfunctional and aggregated proteins as well as damaged organelles. Autophagy thus is a critical catabolic pathway that maintains a functional proteome and sustains cellular integrity and function, while also providing recycled components for anabolism (4–6). Under nutrient-rich conditions, basal autophagy maintains a balanced protein turnover to support growth and biosynthesis. However, under nutrient limitation or other stress conditions, autophagy can be induced to enable metabolic and proteostatic adaptation, thereby maintaining energy homeostasis and survival (5, 7, 8).

Autophagy comprises three mechanistically distinct pathways: macroautophagy (MA), chaperone-mediated autophagy (CMA), and microautophagy. These pathways coexist in mammals, each contributing to the selective turnover of proteins within lysosomes, thereby maintaining cellular quality control (9–12). Macroautophagy has been extensively characterized and is known to play critical roles in stress adaptation, metabolic regulation, and disease processes such as in cancer and neurodegeneration (8, 13–15). It relies on the formation of a double membrane contained autophagosome that fuses with lysosomes for substrate degradation. CMA is specific for soluble proteins containing a targeting motif biochemically related to KFERQ which binds the constitutive chaperone HSC70 (aka HSPA8) to dock on its lysosomal receptor LAMP2A (lysosome associated membrane protein). Upon unfolding, substrate proteins are then translocated by LAMP2A into the lysosomal lumen for degradation (16). Both, MA and CMA can strongly be induced by various forms of cellular stress, often with MA being induced first followed by CMA upon prolonged stress, which is thought to allow a better substrate specificity (17–21).

Endosomal microautophagy (e-MI) is a more recently identified, partially selective autophagic pathway in mammals (22, 23), *Drosophila* (24, 25), and fish (26). During e-MI, cytosolic proteins are directly internalized into late endosomes to form multivesicular bodies (MVBs) for degradation either in the late endosome or lysosomes (22, 24, 25, 27, 28). Unlike macroautophagy, e-MI does not require formation of double-membrane autophagosomes but is characterized by the invagination of endosomal membrane mediated by components of the ESCRT machinery (endosomal sorting complex required for transport). e-MI can occur in bulk or in a substrate specific manner (22, 24, 25, 27, 29–31). Specific substrate recognition in e-MI includes recognition of a KFERQ motif by HSC70 which then binds to phosphatidylserine on the surface of LEs for internalization of the substrate into MVBs (22, 32). Additionally, Hsc70-4 has a membrane bending activity required for e-MI at neuromuscular junctions in *Drosophila*, where it regulates the turnover of synaptic proteins (25). While e-MI in mammals contributes to protein quality control and can process aggregate prone proteins (33), it is mostly constitutively active and not controlled by mTORC1 (28, 31). In *Drosophila*, constitutive e-MI regulates turnover of proteins such as WASp and Comatose/NSF1 (N-ethylmaleimide-Sensitive Factor 1) at neuromuscular junctions in 3^rd^ instar larvae (25). In larval fat body (FB), functionally similar to mammalian liver and adipose tissue (34, 35), e-MI is induced by cellular stress including genotoxic and oxidative stress, and starvation, the latter under negative control by Tor kinase (24, 30). Importantly, as for CMA compared to MA in mammals, stress induction of e-MI requires prolonged starvation: while MA is induced after 1h of starvation and peaks at 4h (36, 37), e-MI takes at least 12 h and peaks at 24 h of starvation (24, 30), suggesting that both processes may degrade potentially distinct sets of target proteins. Since *Drosophila* lack CMA due to absence of a *LAMP2A* homolog in the *Drosophila* genome (18, 38), it has been suggested that *Drosophila* e-MI is an older form of autophagy fulfilling functions that are shared between constitutive e-MI and the starvation-induced CMA in mammals.

Importantly, despite e-MI having been implicated in stress responses and protein quality control (24, 30, 31, 39), little is known about endogenous substrates in *Drosophila*. In this study, we thus used an integrative multi-omics strategy to define the endogenous degradome of e-MI during starvation *in vivo* in *Drosophila* FB, a well-established organ for the study autophagy (24, 30, 34, 36, 37). Our combined proteomic and transcriptomic analyses identified 153 high-confidence e-MI substrates and revealed that ribosomal proteins constituted the most strongly enriched functional category. Starvation induces widespread depletion of proteins involved in translation, aminoacyl-tRNA synthesis, and ribosome biogenesis, whereas the transcript levels of these components remain largely unchanged, indicating significant post-transcriptional regulation. Our study provides the first extensive *in vivo* degradome of e-MI and identifies ribosome turnover as a major physiological function of e-MI in *Drosophila*. Intriguingly, we find distinct translation related changes in FB lacking e-MI or MA, suggesting pathway selectivity that includes distinct subsets of ribosomal proteins. Metabolomic profiling further demonstrated that disruption of e-MI alters starvation-induced metabolic remodeling, highlighting the role of e-MI in coordinating proteome turnover with metabolic adaptation.

## EXPERIMENTAL PROCEDURES

### Fly strains and genetics

Fly food contained 80 g malt extract, 65 g cornmeal, 22 g molasses, 18 g yeast, 9 g agar, 2.3 g methyl para-benzoic acid, and 6.35ml propionic acid per liter. Flies were maintained at a temperature of 25°C with a 12-h light/dark cycle and 50-70% humidity. The following fly stocks were obtained from the Vienna *Drosophila* Stock Center: *Vps32/shrub* RNAi*: KK106823, Vps25* RNAi*: KK108105, Hsc70-4* RNAi: *KK101734, Atg7 RNAi: GD45558. UAS-PA-mCherry-WASp* was from (40)*. cg-Gal4* was a kind gift of Dr. T. Neufeld (University of Minnesota).

### Sample preparation for mass spectrometry

For MS sample preparation, 10-12 virgins flies with appropriate genotype (*cgGal4* was used as wild-type, as it also serves a control for knockdown experiments) were mated to 10 male flies of appropriate genotype (*w^1118^* for wildtype). After 4 days, early third instar larvae were collected from the food and washed twice in water. 50-60 Larvae were then transferred to 35-mm petri dishes for treatment using 3 saturated filter papers (Fisher, #1001–329) soaked in 800µl 20% sucrose supplemented with heat-inactivated yeast (fed) or 20% sucrose only (for starvation) (24). At the end of treatment, larvae were collected and washed twice in water, opened in PBS and inverted with forceps. Fat bodies from approximately 20 larvae were dissected and collected in 50µl RIPA buffer (150mM Nacl, 5mM EDTA, 20mM tween 20; 1% NP40; 0.25% Na-deoxycholate) containing 10mM Benzamidine and homogenized with motorized pestle for 5-10 sec. The samples were centrifuged at 13,000 rpm for 20 mins at 4°C and the supernatant was collected and immediately frozen on dry ice and kept at -20°C. Samples were further processed together using the 96-well S-Trap protocol. 50µl of sample were solubilized in a buffer comprising 5% SDS, 5 mM DTT, and 50 mM ammonium bicarbonate (pH 8) for 1 hour at room temperature to reduce disulfide bonds. The samples were subsequently alkylated with a final concentration of 20 mM iodoacetamide for 30 minutes in the dark. Thereafter, phosphoric acid was added to the sample, achieving a final concentration of 1.2%. Samples were diluted with six volumes of binding buffer (10 mM ammonium bicarbonate, pH 8.0 in 90% methanol). Following gentle mixing, samples were loaded to the S-Trap filter (Protifi) and centrifuged at 500g for 30 seconds. The samples were subsequently washed thrice with binding buffer. Sequencing grade trypsin (1 µg; Promega) diluted in 125µl of 50 mM ammonium bicarbonate (0.008 ug/µl) was then loaded onto the S-Trap column, and the samples were incubated to digestion at 37°C for 18 hours. Peptides were eluted through a three-step process: (1) 80µl of 50 mM ammonium bicarbonate, (2) 80µl of 0.1% trifluoroacetic acid (TFA), and (3) 80µl of 60% acetonitrile with 0.1% TFA. The eluted peptide fractions were subsequently pooled, centrifuged at 1,000g for 30 seconds, and dried using a vacuum centrifuge.

Prior to mass spectrometry analysis, the samples were desalted using a 96-well plate filter (Orochem) containing 1 mg of Oasis HLB C-18 resin (Waters). Briefly, the samples were resuspended in 100µl of 0.1% TFA and applied to the HLB resin, which had been pre-equilibrated with 100µl of the same buffer. Following three washes with 100µl of 0.1% TFA, the samples were eluted using 80 µl of 0.1% TFA and followed by 80 µl of 60% acetonitrile/0.1% TFA. Pooled eluates containing the peptides were dried using a vacuum centrifuge.

### LC MS/MS data acquisition

Samples were resuspended in 10 µl of 0.1% TFA and subsequently loaded onto a Dionex RSLC Ultimate 300 (ThermoFisher), which was coupled online with an Orbitrap Exploris 480 mass spectrometer (ThermoFisher). Chromatographic separation was performed using a two-column system, comprising a C-18 trap cartridge (300 µm inner diameter, 5 mm length) and a picofrit analytical column (75 µm inner diameter, 25 cm length) that was packed in-house with reverse-phase Repro-Sil Pur C18-AQ 3 µm resin (ESI Source Solutions). Peptides were separated utilizing a 90-minute gradient ranging from 4% to 30% buffer B (buffer A: 0.1% formic acid; buffer B: 80% acetonitrile + 0.1% formic acid) at a flow rate of 300 nl/min. The mass spectrometer was configured to acquire spectra in a data-dependent acquisition (DDA) mode. Specifically, the full mass spectrometry scan was set to 300–1,200 m/z in the Orbitrap with a resolution of 120,000 (at 200 m/z) and an AGC (Automatic Gain Control) target of 5 × 10^5^. Tandem mass spectrometry was conducted in the ion trap using the top speed mode (2 s), with an AGC target of 1 × 10^4^ and a higher-energy collisional dissociation energy of 33.

Raw files were searched using Proteome Discoverer software (v2.4, Thermo Scientific) using SEQUEST search engine and the UniProt database of *Drosophila melanogaster*. The comprehensive proteome analysis involved considering N-terminal acetylation as a variable modification and carbamidomethyl cysteine as a fixed modification. Trypsin was designated as the enzyme for digestion, permitting up to two missed cleavages. The mass tolerance was established at 10 ppm for precursor ions and 0.2 Da for-product ions. The false discovery rate for both peptides and proteins were maintained at 1%. All samples were run in quadruplicate for four independent biological replicates.

### MS Data Analysis, bioinformatics and data visualization

Following the search, data was processed as described by Aguilan *et al*. (41). Briefly, proteins were -log_2_ transformed and normalized by subtracting each value from the average value of the respective sample. Missing values were imputed by sampling from a downshifted normal distribution, with the mean shifted by −2 standard deviations relative to the observed intensities, thereby representing low-abundance signals below the detection limit. Statistical significance was assessed using a two-tail heteroscedastic *t*-test (if *p*-value < 0.05). Proteins were ranked using a significance score, which was determined using −log2(p-value) × log_2_(fold change), which integrates statistical significance with direction of change. Data were assumed to be normally distributed.

Bar plots, volcano plots and correlation plots were generated using MS excel. PCA and heatmaps were generated using Perseus (version v2.1.5.0) (42). Log_2_-transformed intensity values were used for visualization, and data were normalized by Z-score transformation across rows. Hierarchical clustering was performed using Euclidean distance with average linkage.

Hub proteins were identified using the CytoHubba plugin in Cytoscape (version 3.10.1) (43) for topological analysis. Nodes were ranked based on maximal clique centrality (MCC) calculated by CytoHubba plugin (44).

Gene Ontology (GO) analysis and biological pathway analysis were performed using go: Profiler (version e113_eg59_p19_6be52918) (45) for different GO molecular function (GO: MF), GO cellular component (GO: CC), GO biological process (GO: BP), and KEGG, and were carried out sequentially for functional annotations. Protein identifiers were mapped to UniProt IDs and analyzed against the *Drosophila* melanogaster reference database (46). Statistical significance was determined using g:SCS multiple testing correction with a significance threshold of 0.05. KEGG Pathway enrichment analysis was performed using Shiny GO 0.77 (47, 48). Enrichment was calculated using a hypergeometric test with Benjamini–Hochberg false discovery rate (FDR) correction for multiple hypothesis testing. The top significant pathways were filtered based on adjusted p value (FDR) < 0.05 and then sorted by Fold Enrichment.

Enriched Reactome pathways were visualized using ReacFoam, a flattened Voronoi representation of the Reactome pathway version 3.7 (49)(https://reactome.org/). Enrichment was assessed by hypergeometric testing against all detected proteins, with p-values corrected using the Benjamini–Hochberg FDR (FDR < 0.05). In this visualization, each polygon represents a pathway, and color intensity reflects enrichment significance (–log_10_ adjusted p-value), with darker shading indicating stronger enrichment.

### RNA-seq

For gene expression analysis, 10-12 *cgGal4* virgins flies were mated to 10 *w^1118^* males. After 4 days, 50-60 early third instar larvae were collected, washed twice in water and transferred to a clean 35-mm petri dishes for 25 h starvation using 3 saturated filter papers (Fisher, 1001–329) soaked in 800 µl 20% sucrose supplemented with heat-inactivated yeast (fed) or 20% sucrose only (for starvation). At the end of 25h, larvae were collected and washed twice in water and inverted in PBS. Fat bodies from approximately 20 fed and starved larvae were collected in biological triplicates, and total RNA extraction was performed using TRIzol reagent according to the instructions by the manufacturer (Invitrogen). 40ng total RNA was sent to Novogene for quality assessment, library preparation, sequencing, and differential expression analysis. Sequencing was performed on an Illumina platform. HISAT2 was used to map the reads to the *Drosophila* genome version 6 (*dm6*) (50). Processing included calculation of the read counts and FPKM (fragments per kilobase of transcript per million base pairs sequenced) values. DESeq2 was used to perform differential expression analysis (51).

### Metabolomics

For metabolome analysis, 10-12 virgins flies with appropriate genotype (*cgGal4* was used for wildtype) were mated to 10 appropriate male flies in fly food containing vials. After 4 days, 50-60 early third instar larvae were washed twice in water and transferred to a clean 35-mm petri dishes for treatment as described above. At the end of 25h, larvae were washed twice in water and inverted in PBS. ∼35 mg fat bodies from 25-30 fed and starved 3^rd^ instar larvae of indicated genotypes for each of six biological replicates were collected in 100µl ice cold PBS in a 2ml screw cap tube (USA scientific #1420-8799). PBS was removed and fat bodies were washed with 100µl of 150mM ammonium acetate and centrifuged at 1,200 rpm for 10 mins at 4 ^0^C. The buffer was removed and fat bodies were snap frozen on dry ice and stored at -80 ^0^C until use. Samples were extracted in 200µl of methanol:acetontrile:water = 2:2:1 and internal standards were added to each sample (Supplementary table 1). After vortexing, the samples were lysed by three freeze-thaw cycles in liquid nitrogen followed by ice-cold sonication. Samples were then centrifuged at 14000rpm for 10 min and the supernatant was transferred to glass vials for analysis by the Einstein Metabolomics facility.

Briefly, the samples were analyzed with ABsciex 6500+ MS with an iHILIC-p column (HILICON) and an Ace PFP column. A pooled quality control (QC) sample was included in the sample list and injected six times to calculate the coefficient of variation (CV) for data quality control. Metabolites with CV less than 30% in the QC samples were used for quantification after combining the results from both columns. For quantification, data were -log_2_ transformed and normalized by subtracting each value from the average value of the respective sample. Missing values were imputed by sampling from a downshifted normal distribution, with the mean shifted by −2 standard deviations relative to the observed intensities, thereby representing low-abundance signals below the detection limit. Statistical significance was assessed using a two-tail heteroscedastic *t*-test (if *p*-value < 0.05). Metabolites were ranked using a significance score, which was determined using −log_2_(p-value) × (log_2_ fold change), which integrates statistical significance with direction of change. Bar plots and volcano plots were generated using MS excel. MetaboAnalyst 6.0 (https://www.metaboanalyst.ca/) (52) was used to generate PLS-DA score plot.

### Joint pathway enrichment analysis

For an integrative omics-data analysis, wild type significantly altered proteins and metabolites (FDR ≤ 0.05) were submitted to MetaboAnalyst 6.0 (https://www.metaboanalyst.ca/) (52). Protein names and HMDB identifiers were submitted alongside their log_2_-fold changes between starved vs fed condition. For the Joint Pathway Enrichment Analysis, the Gene-Metabolite-Interaction network was selected based on the organism *Drosophila melanogaster* and the KEGG reference pathway database. Default settings were chosen with a hypergeometric test and the topology measure with the degree centrality. The applied integration method was the default option “combine queries”. The analysis was conducted for metabolic integrated pathways as well as all integrated pathways.

### e-MI assays and Image analysis

e-MI assays were performed as described in Mukherjee et al. (24). Briefly, 10-12 *UAS-KFERQ-PA-mCherry; cgGal4* virgins were mated to 10 males of appropriate genotype (*w^1118^* for controls) in fly food containing vials. After 4 days, 50-60 early third instar larvae were collected from the food, washed twice in water, and transferred to a clean 35-mm petri dishes containing 800 µl Graces insect medium (Invitrogen, 11605-094) with 10% heat-inactivated fetal bovine serum (Atlanta Biochemicals, S11050). The KFERQ-PA-mCherry sensor was photoactivated by exposure to a 405-nm light source for 11 min (at 2.8 A, approximately 60 µW/cm2) (24). Immediately after photoactivation, larvae were washed twice in water and transferred to a clean 35-treated for starvation for 25 h using 35-mm petri dishes with 3 saturated filter papers (Fisher, 1001–329) soaked in 800 µl 20% sucrose supplemented with heat-inactivated yeast (fed) or 20% sucrose only (for starvation). Larvae were washed twice in water and inverted in PBS and fixed in 4% paraformaldehyde in phosphate-buffered saline (PBS) for 1 h at room temperature (RT). Samples were then washed with PBS for three 15 min cycles at RT. Fat bodies from 5-10 larvae were then dissected and mounted in 20µl DAPI Fluoromount-G (SouthernBiotech, OB 0100-20) per slide and stored overnight at 4°C prior to imaging. *Drosophila* fat bodies expressing KFERQ biosensor were imaged using a 63 × 1.4 NA oil objective on an ApoTome.2 system (Carl-Zeiss, Oberkochen, Germany). Quantification of puncta was done using SParQ plug-in in ImageJ/Fiji (53).

### Statistical analysis

Statistical analyses were done using GraphPad Prism (Versions 8 and 9; GraphPad Software, La Jolla, CA) with the indicated tests and corrections. Unless otherwise noted, graphs depict means with standard error of the means (SEM). The notation “n=” refers to fields of view unless specified otherwise. Given that larvae exhibit bilateral symmetry and possess two primary FB lobes, it is possible that two fragments may have originated from the same lobe during preparation. Consequently, it is a reasonable estimation that the number of animals is at least approximately n/2 and at maximally equal to n.

## RESULTS

### Starvation induces significant changes in the Drosophila fat body proteome

To date, only very limited information is available regarding physiological substrates of *Drosophila* e-MI, and no systematic approach has been undertaken in any organ for their identification (25, 54). We thus started to define the consequences of prolonged nutrient deprivation known to induce a strong e-MI response in the *Drosophila* larval FB and performed quantitative LC MS/MS proteomic analyses on the fat bodies of wild-type (WT) third-instar larvae starved for 25h on 20% sucrose (Fig. 1A). Principal component and hierarchical clustering analyses revealed clear segregation between fed and starved samples, indicating a robust and reproducible starvation-induced proteomic response (Fig. 1B; Fig. S1A, B; note that our final analysis focused on three replicates of starved FB). Differential abundance analysis identified 160 proteins significantly increased or decreased in abundance in response to prolonged starvation in WT (p*-*value < 0.05 or >4.32 when -log_2_ transformed; Fig. 1C; Supplementary table 2).

**Figure 1:**
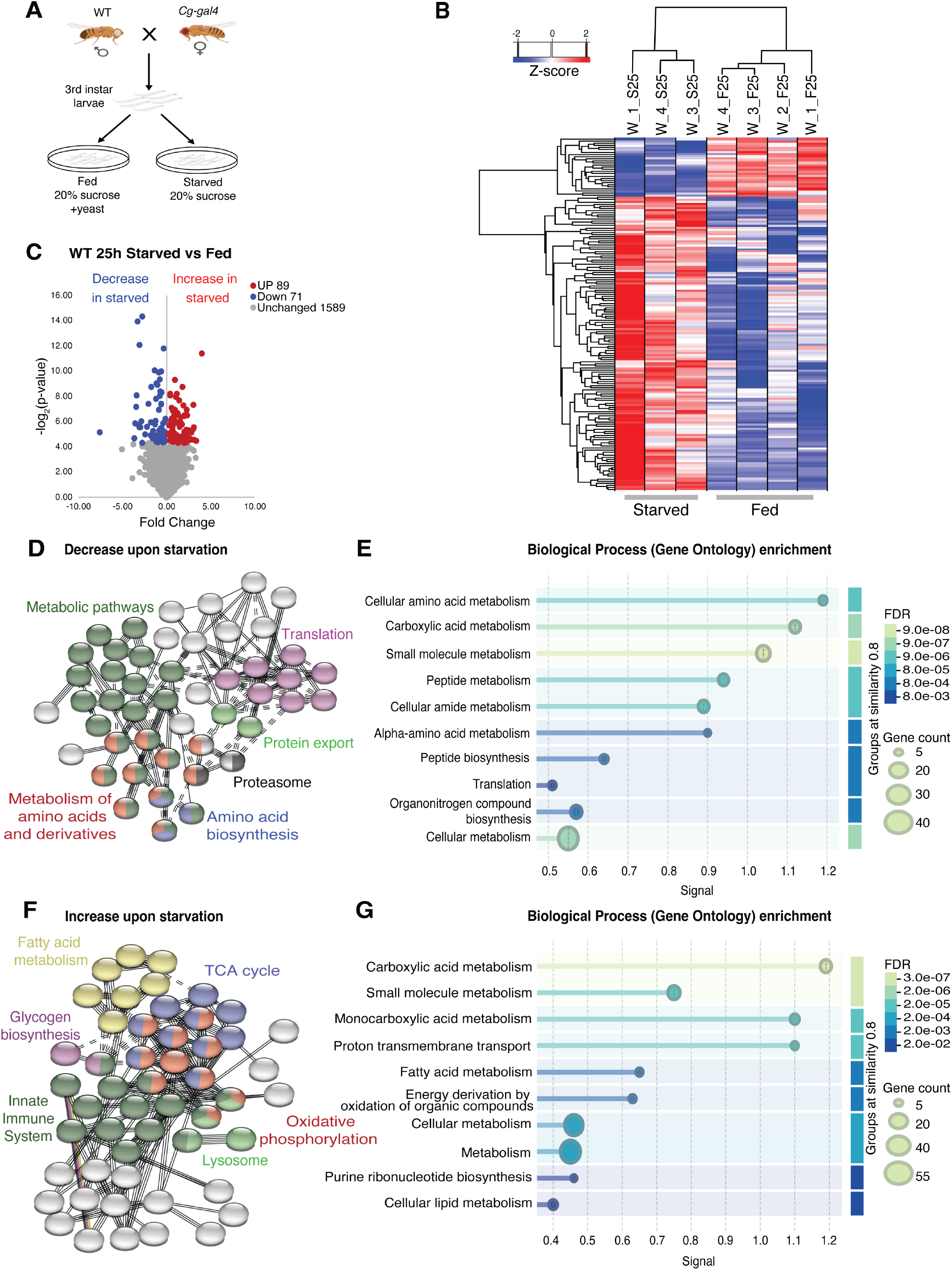
The *Drosophila* fat body proteome in response to prolonged starvation. **(A)** Experimental overview of e-MI induction by starvation in *Drosophila* larvae. FB tissues were dissected and analyzed by quantitative LC–MS/MS. **(B)** Heatmap of proteins with significantly changed abundance (p-value>3 when -log_2_ transformed; red = stabilized, blue = degraded) across fed and starved samples highlighting the global proteomic differences between the conditions. **(C)** Volcano plot of fold changes in protein abundance (starved/fed) showing 160 significantly altered proteins. p-value>4.32 when log2-transformed (red = up, blue = down). **(D)** STRING protein–protein interaction network of proteins decreased upon starvation revealed clusters involved in translation, protein export, proteasome, amino acid biosynthesis, metabolism of amino acids and derivatives, and metabolic pathways. **(E)** The corresponding top 10 GO enrichment terms using STRING database showed enrichment of translation, peptide biosynthesis, and amino acid metabolism. **(F)** STRING interaction network of proteins increased under starvation. **(G).** The corresponding top 10 GO enrichment analysis using string database for biological processes. In (E, G) the x-axis indicates enrichment significance (-log₁₀ FDR) and the y-axis lists GO biological process terms. Bubble size reflects the number of proteins associated with each term, and color intensity corresponds to the false discovery rate (FDR) as per legends to the right.

To identify functionally connected proteins within the starvation-downregulated proteome, we performed STRING interaction analysis and Gene Ontology enrichment of starvation-downregulated proteins which showed significant enrichment of translation, peptide biosynthesis, and amino acid metabolism (Fig. 1D,E). To further define key regulators (hub proteins) within this protein-protein interaction network, we performed centrality analysis using the CytoHubba plugin in Cytoscape and ranked nodes by maximal clique centrality (MCC) to highlight a core set of hub proteins that were preferentially reduced upon starvation (Fig. S1C). The top 10 hub proteins include components of the translational machinery, such as ribosomal proteins (Rpl20-like, Rpl9), translation initiation and termination factors (eIF2α, eIF3m, eRF1), and aminoacyl-tRNA synthetase (ArgRS). Additional hub proteins are associated with ribosome biogenesis (CG5728/Rrp5, CG9246/Noc2 protein), ribosome associated stress sensing (l(3)80fj, ortholog of human GCN1 protein), and co-translational targeting to ER (SrpRα), which suggests the coordinated regulation of ribosome homeostasis and translation in response to starvation. Together, these findings indicate that starvation-induced proteome remodeling preferentially targets ribosome related and translational machinery.

Next, we performed centrality analysis of proteins significantly increased upon starvation (Fig. S1D). MCC ranking highlighted hub proteins associated with mitochondrial and energy metabolism, including ATP synthase subunits (ATPsynB, ATPsynF, ATPsynγ), electron transport chain components (ND-20, UQCR-Q), and vesicular acidification machinery (VhaAC39-1, Vha100-2). Gene ontology enrichment analysis of proteins significantly increased upon starvation demonstrated significant enrichment of oxidative phosphorylation, lipid and fatty acid metabolism, and energy generation pathways (Fig. 1F,G). Collectively, these data indicate that starvation, which here primarily reflects removal of yeast as a protein source while maintaining sucrose as a carbon source, promotes coordinated upregulation of mitochondrial and metabolic networks alongside selective suppression of translational capacity. This distinct regulation suggests that starvation induces the reallocation of cellular energy resources with suppression of protein synthesis and activation of mitochondrial pathways that support energy production.

To determine whether starvation-induced proteomic remodeling is transcriptionally driven, we analyzed transcriptional changes that occur in the FB upon prolonged starvation and integrated our proteomic with matched transcriptomic data. We found that the mRNA levels of the vast majority of genes (10600) were unaffected by prolonged starvation of 25h while 1994 and 1190 genes were transcriptionally induced or down regulated, respectively (Fig. 2A; Supplementary table 3). Many proteins that decrease in abundance upon starvation do so without corresponding reductions in mRNA levels (highlighted by orange data points in Fig. 2B; Supplementary table 3) suggesting that their downregulation is largely driven by increased protein degradation or less synthesis.

**Figure 2.**
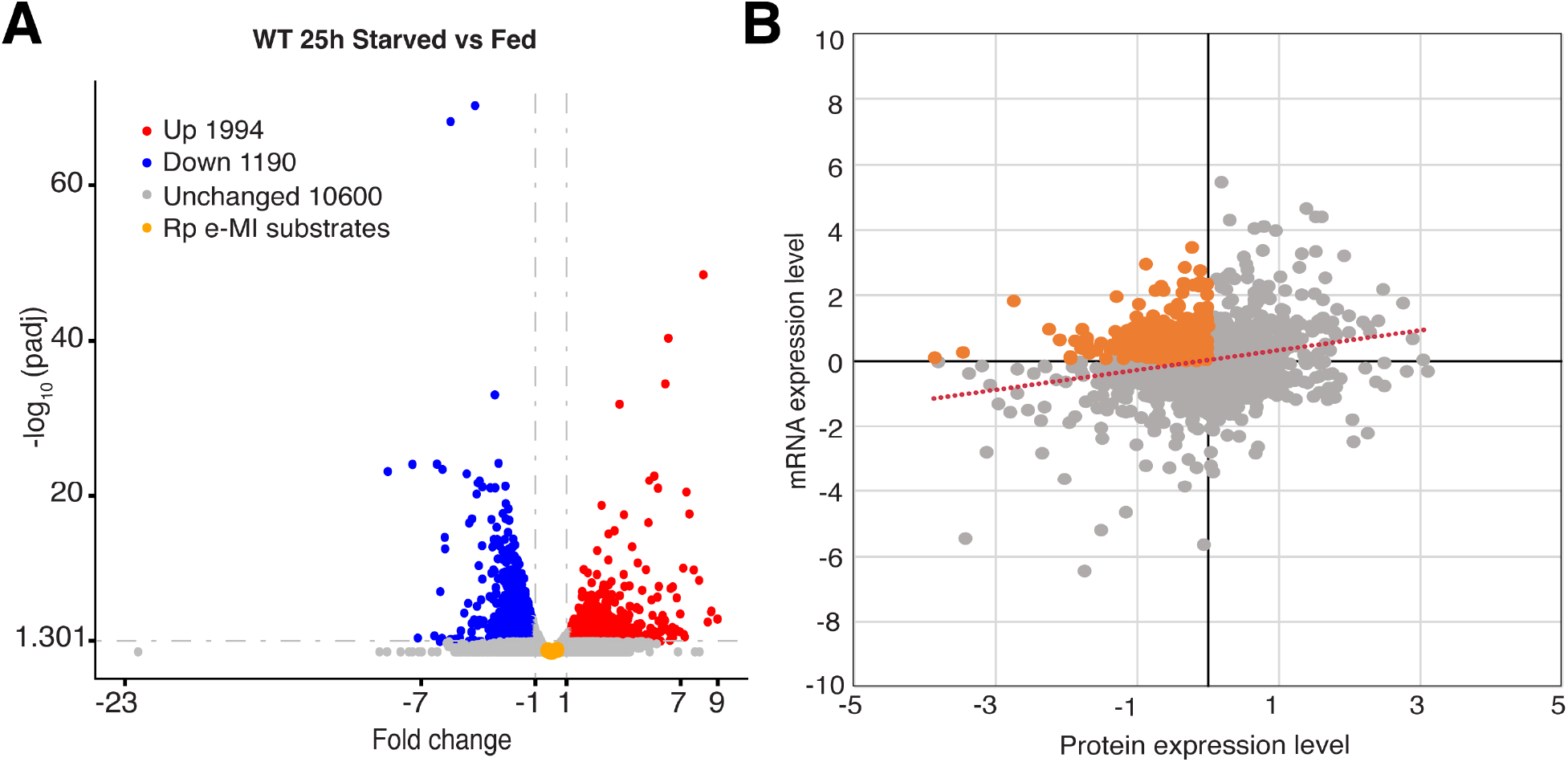
Transcriptomic changes due to starvation in the fat body. (**A**) Volcano plot of differentially expressed genes in WT FB upon 25 h starvation. Cutoffs were adjusted p <0.05 and log_2_ fold change ≥ 1. Red, blue, and grey circles depict up-, down-regulated, and unchanged genes, respectively. ‘Red’ and ‘Blue’ genes were used to filter proteome analysis. Orange dots represent ribosomal proteins that are e-MI substrates. (**B**) Correlation of fold changes upon 25h starvation between matched WT 25h starved vs fed transcript and WT 25h starved vs fed protein levels. Each point represents a gene–protein pair. Orange points indicate increased/unchanged mRNA but low protein abundance upon starvation, consistent with post-transcriptional regulation. Red line represents linear regression fit to assess correlation between the two datasets.

### Identification of endosomal microautophagy substrates in the fat body

e-MI depends on MVB formation by the ESCRT machinery and -at least in part-on Hsc70-4 (24, 25). Endogenous e-MI substrates are thus expected to be stabilized in the absence of ESCRT components or Hsc70-4. Similarly, we recently found that overexpression of the actin nucleation factor WASp prematurely induces e-MI after 4 h of starvation without changing MA (40) and it is thus expected that WASp overexpression will induce degradation of e-MI substrates at this time point. e-MI activity in the fly FB can be measured using a photoactivatable KFERQ-PAmCherry reporter that relocates to (endo)lysosomes upon starvation (Fig. 3A,B and F, G) (24). RNAi-mediated depletion of the ESCRT III component *Vps32* or the *Hsc70-4* in the FB using the UAS-Gal4 system (55) significantly reduced starvation-induced e-MI reporter puncta (Fig. 3,D and I; quantified in Fig. 3E, J), demonstrating efficient *in vivo* knockdown. We thus investigated whether e-MI contributes to starvation-induced proteome remodeling using the above proteomic approach and compared protein abundance profiles of wild-type fat body with those of FB-specific *Vps32* RNAi and *Hsc70-4* RNAi larvae following 25h of starvation (Fig. S2; Supplementary table 2). Differential abundance analysis revealed widespread alterations in protein levels upon their knockdown, with proteins enriched in WT (positive fold change, red) and a distinct set enriched in the corresponding RNAi condition (negative fold change, blue). In *Vps32* RNAi FB, proteins significantly accumulating were linked to endomembrane trafficking and cellular stress responses, including ESCRT component TSG101, stress responsive kinases p38 and the ER stress associated Hacl protein indicating altered cellular trafficking and proteostasis. Additionally, Rhea and Inos proteins involved in membrane organization were also higher in *Vps32* RNAi FBs. On the other hand, proteins significantly higher in wildtype were predominantly components of mitochondrial and metabolic pathways such as oxidative phosphorylation subunit ND75, ATPase synthetase subunit ATPsynB and mitochondrial ribosomal protein mRpL37, as well as metabolic enzymes such as CTPsyn and Taldo, indicating enrichment of mitochondrial and oxidative metabolism (Fig. S2A). KEGG pathway enrichment analysis also revealed distinct responses to prolonged starvation of 25 h in WT and ESCRT-depleted fat bodies (Fig. S2B). ESCRT RNAi samples were enriched for amino acid metabolism (including arginine biosynthesis, alanine, aspartate and glutamate metabolism, tryptophan metabolism and branched-chain amino acid degradation), fatty acid degradation, TCA cycle, glutathione metabolism and proteasome and lysosome indicating metabolic stress. In contrast, wild-type showed enrichment of lipid and mitochondrial metabolism, including fatty acid metabolism, the TCA cycle, oxidative phosphorylation and pyruvate metabolism. These were accompanied by broad amino acid metabolic pathway changes and aminoacyl-tRNA biosynthesis, indicating coordinated remodeling of energy production, biosynthetic capacity and reliance on oxidative metabolism. Together, these findings indicate that loss of Vps32 function disrupts normal metabolic adaptation to starvation and instead promotes stress associated metabolic state, highlighting a requirement for ESCRT in maintaining metabolic homeostasis.

**Figure 3:**
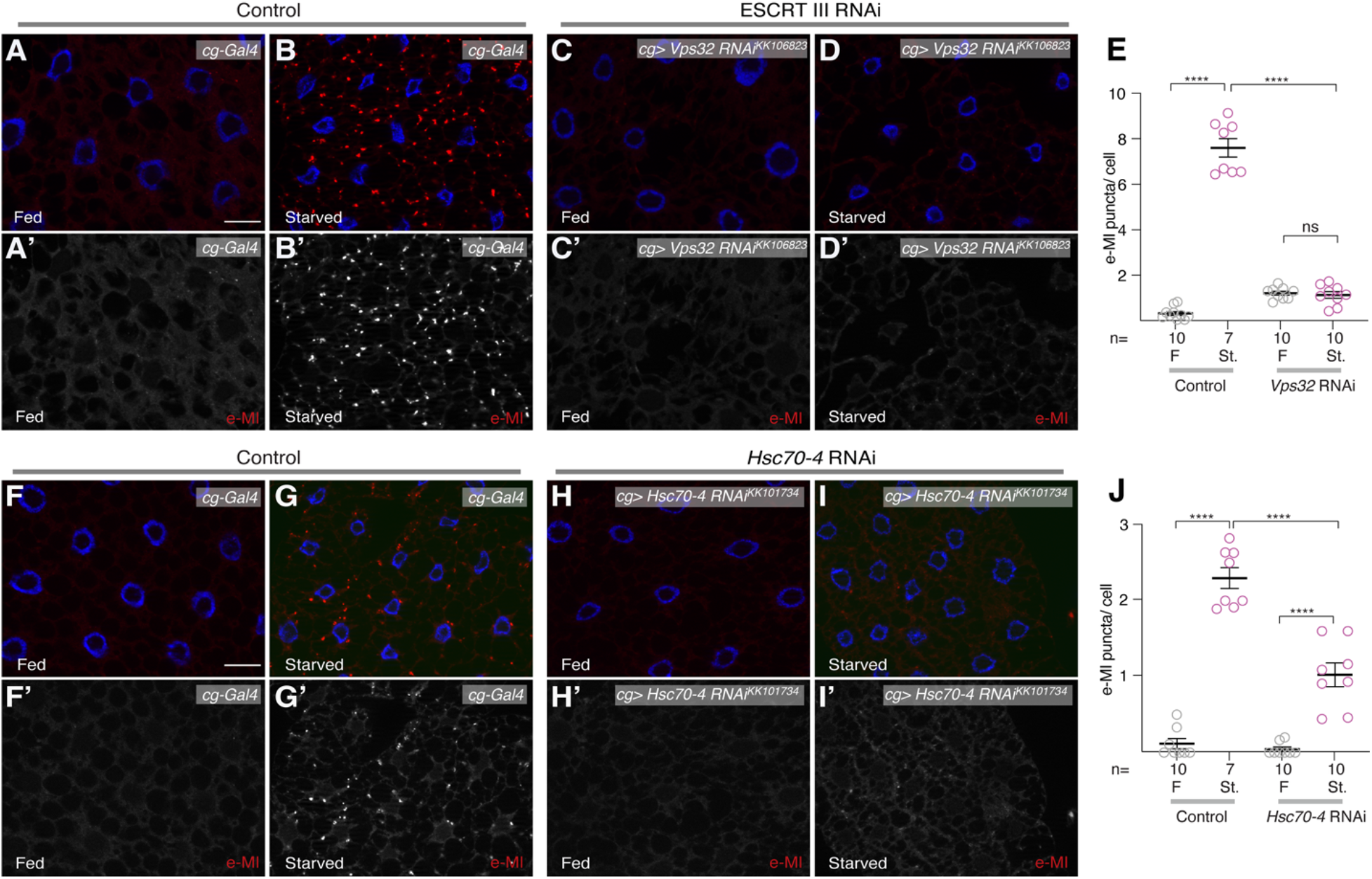
RNAi mediated knockdown of the *ESCRT III* component *Vps32* (*shrub*) and *Hsc70-4* inhibits starvation induced e-MI. (A-D and F-I) Representative images of 3^rd^ instar larval FB showing e-MI activity (KFERQ-PA-mCherry reporter puncta) in control (A,B and F,G), *Vps32* (C,D), and *Hsc70-4* (H,I) *RNAi* larvae. While starvation induces e-MI in control (red puncta in B, G), knockdown of *Vps32* and *Hsc70-4* strongly reduces the e-MI response (D, I). Greyscale images show e-MI activity; nuclei are in blue; scale bar 20µm. **(E, J)** Quantification of e-MI response as puncta per cell at 25 h of treatment. Data are mean ± SD. One way ANOVA p<0.0001 (Tukey); \*\*\*\**p* < 0.0001, ns: not significant.

Hsc70-4 RNAi causes a significant accumulation of translation and ribosome related proteins (e.g. eIF3m, Bys and mRpl35) along with the metabolic proteins such as NT5E-2, stress response proteins CkIIβ and cytoskeletal organization proteins Rhea and Tm2 which suggests impaired turnover of ribosomal and cytosolic components (Fig. S2C). It also caused a decrease of factors involved in protein folding such as Hsp3 and CCT1, energy metabolism (Had1, Adh, SesB) and cell cycle regulation protein Ran. KEGG pathway analysis of proteins increased in *Hsc70-4* RNAi indicated strong enrichment for multiple metabolic pathways of amino acid and intermediary metabolism, including alanine, aspartate and glutamate metabolism, beta-alanine, propanoate, glycine, serine and threonine metabolism, as well as pyrimidine and purine metabolism. Additional enrichment was observed for proteasome, pentose phosphate pathway, oxidative phosphorylation, glutathione metabolism, biosynthesis of amino acids, and carbon metabolism pathways. KEGG analysis of *Hsc70-4* RNAi decreased proteins identified a broad set of metabolic pathways, including glyoxylate and dicarboxylate metabolism, TCA cycle, pyruvate metabolism, glycolysis/gluconeogenesis, fatty acid degradation, and multiple amino acid metabolism pathways (including valine, leucine and isoleucine, glycine, serine and threonine; Fig. S2D) indicating depletion of Hsc70-4 is associated with decreased central carbon metabolism, energy production and biosynthetic adaptation pathways during starvation(56, 57).

To assess how premature activation of e-MI remodels the proteome, we compared wild-type larval FB with FB overexpressing WASp after 4h of starvation. Proteins involved in metabolic and stress response pathways such as Adk2, HisRS, and Nrg significantly decreased at 4 h starvation upon Wasp overexpression (Fig. S2E), while factors associated with translation and cellular homeostasis, such as eIF2β, eIF2-RB, NimB2, and SERCA increase. The data indicate that while the majority of the proteome remains relatively stable between conditions, a defined subset of proteins shows differential abundance between UAS-WASp and WT samples. (Fig. S2E; Supplementary table 2), mirroring some of the patterns observed at later time points in wild-type starvation. KEGG pathway enrichment analysis suggests that WASp expression enriched for pathways associated predominantly with central carbon metabolism, including glycolysis/gluconeogenesis, the citrate cycle, carbohydrate metabolism and amino acid metabolism (Fig. S2F). In contrast, proteins stabilized under those conditions are enriched for sulfur metabolism, lipid (fatty acid biosynthesis, fatty acid degradation, arachidonic acid metabolism), and amino acid metabolism, as well as pathways involved in detoxification such as drug and cytochrome P450-mediated metabolism (Fig. S2F).

The proteins reduced upon early e-MI activation by overexpression of Wasp showed functional overlap with proteins that are significantly decreased during prolonged starvation in wild-type FB at the level of enriched KEGG pathway. We found consistent enrichment of pathways involved in amino acid metabolism, organic/carboxylic acid metabolism and central carbon metabolism.

### e-MI targets ribosomal and translation associated proteins

We then combined the various genotypes and conditions according to the anticipated fate of true e-MI substrates and implemented an integrated workflow combining proteomics and RNA-seq with the genetic perturbations of the e-MI machinery, and temporal starvation paradigms (Fig. S3). Under physiological conditions, e-MI substrates are expected to be degraded upon prolonged starvation (25 h; Fig S3). Similarly, they should be prematurely degraded upon overexpression of WASp already after 4 h of starvation (40). We thus combined the starved vs fed ratios of proteins identified in our mass spec data of those two situations as our baseline for comparison. Proteins that decreased during starvation in wild type and UAS-WASp backgrounds but failed to decrease in *Vps32* and/or *Hsc70-4* RNAi conditions (Fig S3) were then identified by comparison with the starved vs fed ratios of proteins identified under those conditions. At the same time, we only focused on proteins the mRNA of which either did not change or upregulated upon starvation.

This approach led to the identification of 153 high scoring e-MI candidate substrates (Supplementary table 2), functional classification of which revealed striking enrichment for ribosomal proteins, translation-related components, and lysosomal proteins (Fig. 4). Particularly, GO enrichment analysis identified ‘structural constituents of ribosomes’ (molecular function), ‘cytoplasmic TL’ (biological process), ‘cytosolic ribosome’, and ‘ribosomal subunit’ (cellular component) as top enriched terms respectively (Fig. 4A). The top enriched pathway in the KEGG analysis also is ‘ribosome’ (Fig. 4A). Moreover, KEGG pathway analysis using ShinyGO also shows significant enrichment of the ‘ribosome’ and ’lysosome’ (Fig. 4B; FDR = 1.4×10⁻^4^ and 2.2×10⁻3, respectively). The effect on ribosomal proteins is also clearly seen in a heatmap of all significantly changed ribosomal protein levels averaged across all biological replicates for the different genotypes showing clear stabilization upon disruption of e-MI machinery (Fig. 4C). String network analysis showed the expected dense connectivity among both large and small ribosomal subunits (Fig. 4D), while RNA-seq data indeed confirmed that corresponding transcripts were not significantly changed (Fig. 2A, orange dots). GO enrichment analysis further highlighted translation, peptide biosynthesis, and organonitrogen compound biosynthesis as the most significantly enriched biological processes among candidate e-MI substrates (Fig. 4F). Additionally, Reactome pathway enrichment analysis identified translation-related pathways within the “metabolism of proteins” category as significantly enriched, with strong representation of eukaryotic translation initiation, SRP-dependent co-translational protein targeting to membrane, and ribosome-associated quality control (Fig. 4G). Together, these data reveal that ribosomal proteins constitute a major class of e-MI substrates during starvation.

**Figure 4:**
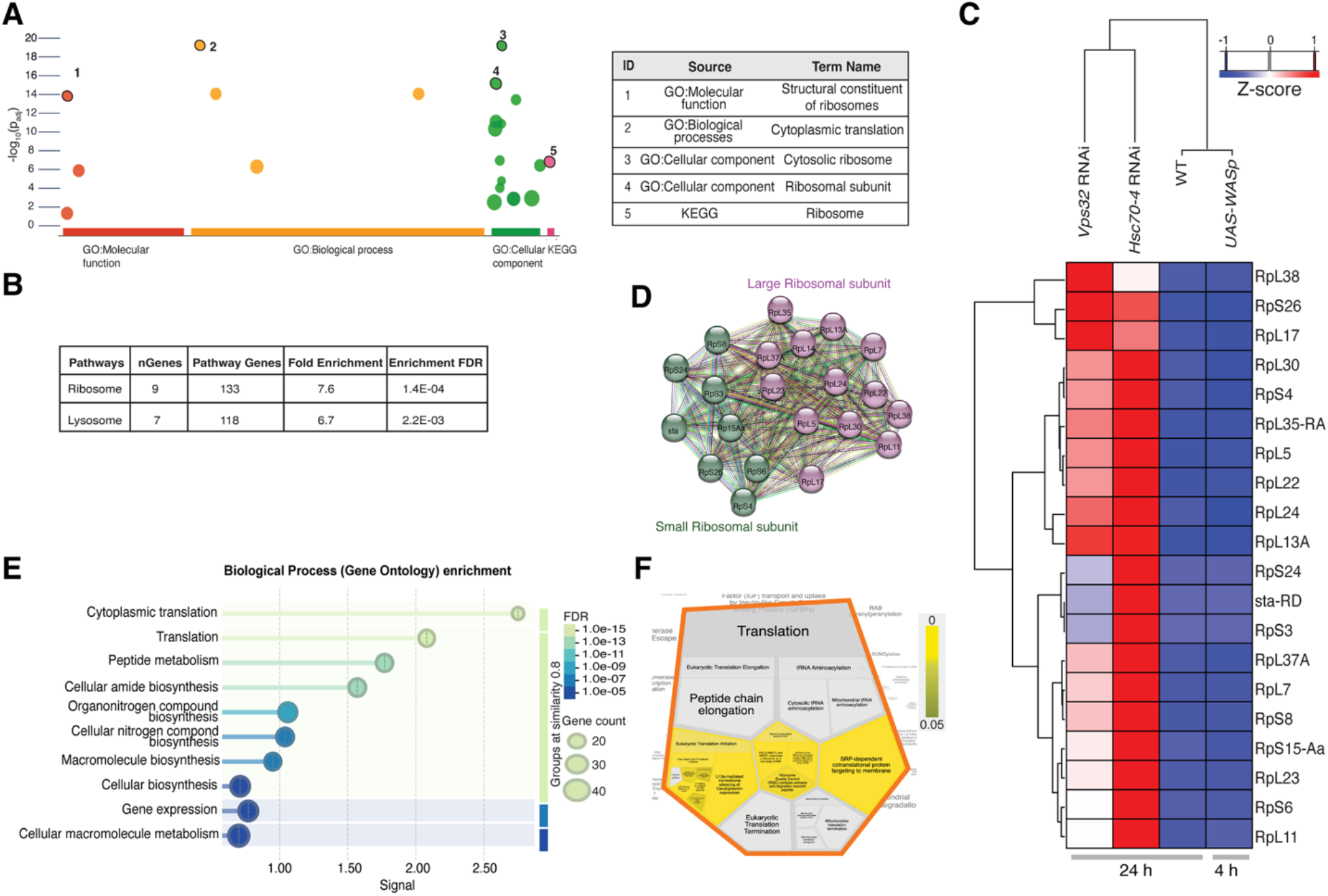
Ribosomal proteins are a major class of e-MI substrates. **(A**) Top term hits identified upon functional enrichment analysis of significant e-MI substrate using g:Profiler. Enriched terms are plotted by statistical significance with circle size reflecting the number of proteins associated with each term. Numbered terms correspond to the entries listed in the table on the right. **(B)** KEGG pathway enrichment analysis of 153 candidate substrates using ShinyGO revealed ‘ribosome’ and ‘lysosome’ as the strongest enriched pathways. **(C)** Heatmap of fold changes averaged over biological replicates of all significantly changed ribosomal protein across different genotypes under starvation in WT (25 h; n = 3), *UAS-WASp* (4 h ; n = 4), *Vps32* RNAi (25 h; n = 4), and *Hsc70-4* RNAi (24 ;n = 4). Ribosomal proteins are depleted in WT and *UAS-WASp* backgrounds (in blue) but stabilized upon disruption of endosomal microautophagy machinery (in red). **(D)** STRING interaction network of ribosomal proteins identified as e-MI substrates. **(E)** Top 10 GO terms of candidate e-MI substrates for biological process using STRING database showed cytoplasmic translation and translation as the most enriched pathways. The x-axis indicates enrichment significance (-log₁₀(FDR)) and the y-axis listing GO biological process terms. Bubble size reflects the number of proteins associated with each term, and color intensity corresponds to the false discovery rate (legends on the right). **(F)** The Reactome pathway enrichment map of all e-MI candidate substrates highlights translation initiation, elongation, termination, tRNA aminoacylation, and SRP-dependent membrane targeting pathways as significant subgroups Tile size reflects gene counts and color reflects enrichment significance.

### e-MI and macroautophagy target distinct proteins

These findings prompted us to also assess changes in the proteome upon inhibition of MA, peaking around 4 h of starvation (37). We first verified that autophagosome formation is blocked by RNAi mediated knockdown *Atg7*, a gene essential for autophagosome initiation in the larval FB (58). In control larvae, short-term starvation (4 h) induced robust accumulation of autophagosomes as assessed by mCherry–Atg8a puncta formation (*cg-Gal4*; Fig. 5A,B). RNAi-mediated knockdown of *Atg7* effectively suppressed starvation-induced Atg8a puncta formation, confirming efficient inhibition of macroautophagy under these conditions (Fig. 5C,D; quantified in E). Comparing the proteome of WT and Atg7 knockdown FB following 4 h starvation revealed that 272 proteins were stabilized upon inhibition of MA, while 304 were of lower abundance (Fig. 5F; Supplementary table 2). Comparative KEGG pathway enrichment analysis showed distinct pathways affected in WT and *Atg7* RNAi 4 h post starvation (Fig. 5G). Enriched pathways identified among candidate MA substrates are metabolism of amino acids, fatty acid degradation, peroxisome, along with oxidative phosphorylation and ER protein processing. *Atg7* RNAi led to the increase of pathways related to the central carbon metabolism including pentose phosphate pathway, glycolysis, TCA cycle and biosynthetic processes such as, amino acid biosynthesis, lipid metabolism, and redox pathway (Fig. 5G). Together, these results suggest that, during starvation, WT tissues undergo a coordinated metabolic adaptation facilitated by Atg7-dependent autophagy, which supports sustained energy production, biosynthesis, and redox homeostasis. Thus, Atg7-dependent autophagy required not only for degradation but also for maintaining metabolic balance and organelle homeostasis during starvation, while loss of Atg7 results in a shift to metabolic imbalance and cellular stress.

**Figure 5:**
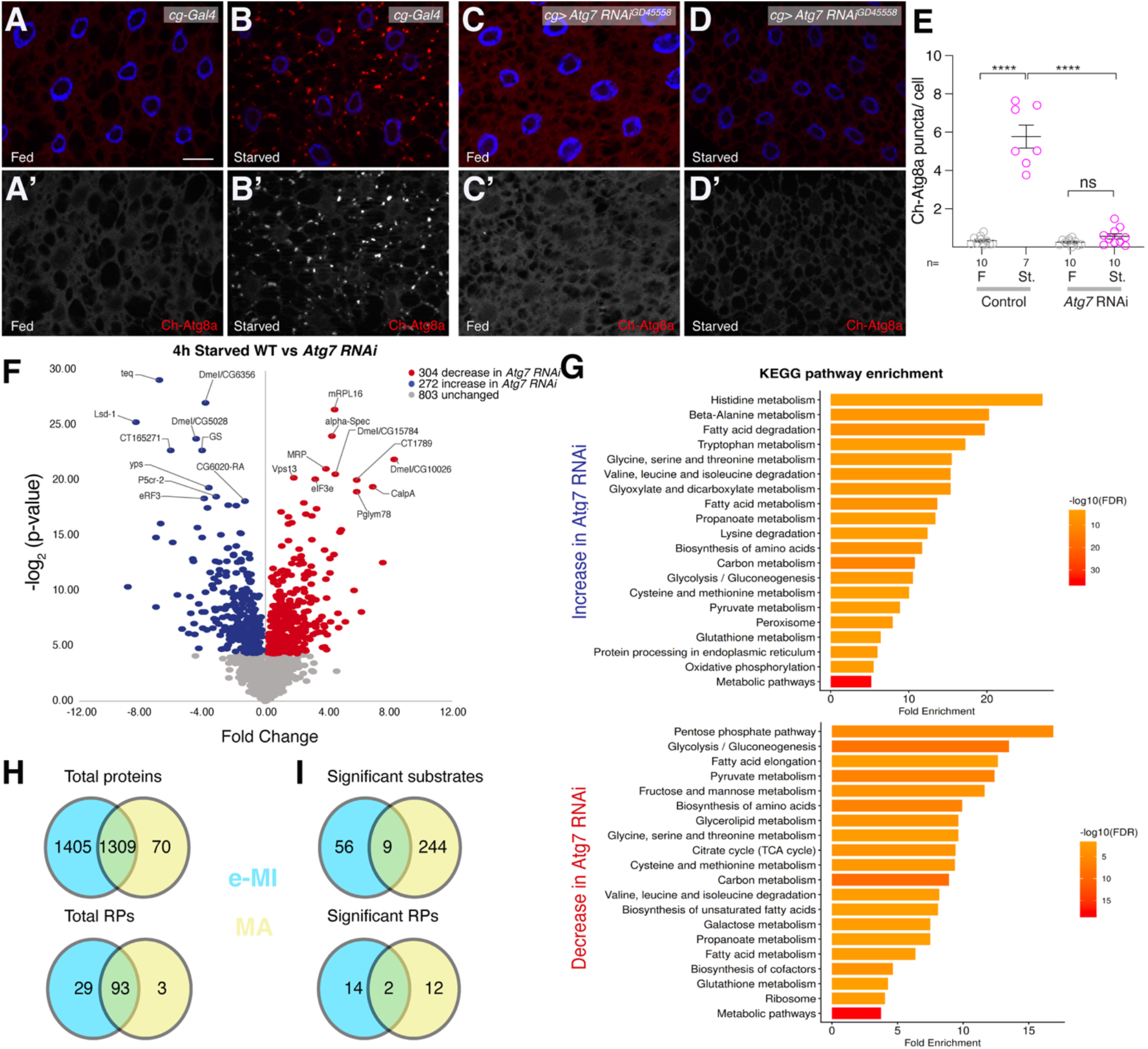
Identification of candidate macroautophagy substrates. (A-D) Compared to fed control (A), 4h starvation robustly induces MA as assessed by the formation of mCherry-Atg8a puncta marking autophago(lyso)somes (B). MA is prevented by RNAi mediated knockdown of *Atg7^GD45558^* (C, D). Nuclei are stained with DAPI (blue); greyscale panels show mCherry-Atg8a. Scale bar: 20µm. **(E)** Quantification of mCherry-Atg8a puncta in indicated conditions and genotypes. One way ANOVA p<0.0001 (Tukey); **** *p*<0.0001, ns: not significant. **(F)** Volcano plot showing differential protein abundances in 4h starved WT vs *Atg7* RNAi *FB.* Candidate MA substrates are in blue, proteins that are increased upon Atg7 RNAi knockdown are in red (*p*-value cutoff <0.05; >4.32 when -log_2_ transformed). **(G)** Bar plot showing KEGG pathway enrichment analysis of proteins that are significantly higher in *Atg7* RNAi (top) and decrease in *Atg7* RNAi (bottom) 4h post starvation using the ShinyGO database. The x-axis indicates ranked fold enrichment. Bar color represents statistical significance as -log_10_(FDR), with darker colors indicating higher significance (scale on right). **(H)** Venn diagrams showing the overlap of total proteins (above) and ribosomal proteins (below) identified in the e-MI (blue) and MA (yellow) MS experiments. **(I)** Among the 1309 proteins shared in both datasets, e-MI (blue) and MA (yellow) substrates are largely distinct (9 shared; above). Among the substrates, most ribosomal proteins (below) are also pathway-selective substrates.

We then addressed if e-MI and MA selectively degrade proteins. To do so, we first identified all proteins that were called by MS in both, the e-MI and MA parts of our study, which resulted in 1309 proteins (Fig. 5H; these included 93 cytoplasmic and mitochondrial ribosomal proteins). Interestingly, among those, the identified e-MI and MA substrates seem to be largely distinct. Of the 65 likely e-MI and 243 MA substrates, only 9 are shared suggesting an unanticipated extent of autophagic pathway specificity (Fig. 5I). Importantly, even among the 16 and 14 ribosomal proteins identified as eMI and MA substrates, respectively, only two are shared (Fig. 5I), supporting the conclusion that Atg7-dependent macroautophagy and e-MI contribute selectively to proteome regulation during starvation.

### Metabolomic profiling reveals e-MI-dependent metabolic remodeling during starvation

As our proteome analyses indicated that metabolism including the one of amino acids may be affected by prolonged starvation, we performed untargeted metabolomics on fat bodies from wild-type, *Vps25 (ESCRTII) RNAi*, and *Hsc70-4* RNAi larvae under fed and starved conditions (25h; Fig. 6). Principal component analysis showed clear separation between genotypes, indicating distinct metabolic responses (Fig. 6A). In wild-type animals, starvation induced a reduction across multiple metabolite classes, including essential and non-essential amino acid (blue and red, respectively in Fig. 6B; Supplementary table 4). In contrast, inhibition of eMI by *Vps25* (ESCRT II) or *Hsc70-4* knockdown markedly altered this amino acid metabolic response (Fig. 6 B,E; Supplementary table 4). The amino acid depletion is attenuated upon prolonged starvation, with essential amino acids exhibiting smaller changes, indicating tighter regulation of their abundance during starvation (Fig. 6 C,D,F,G; Supplementary table 4).

**Figure 6:**
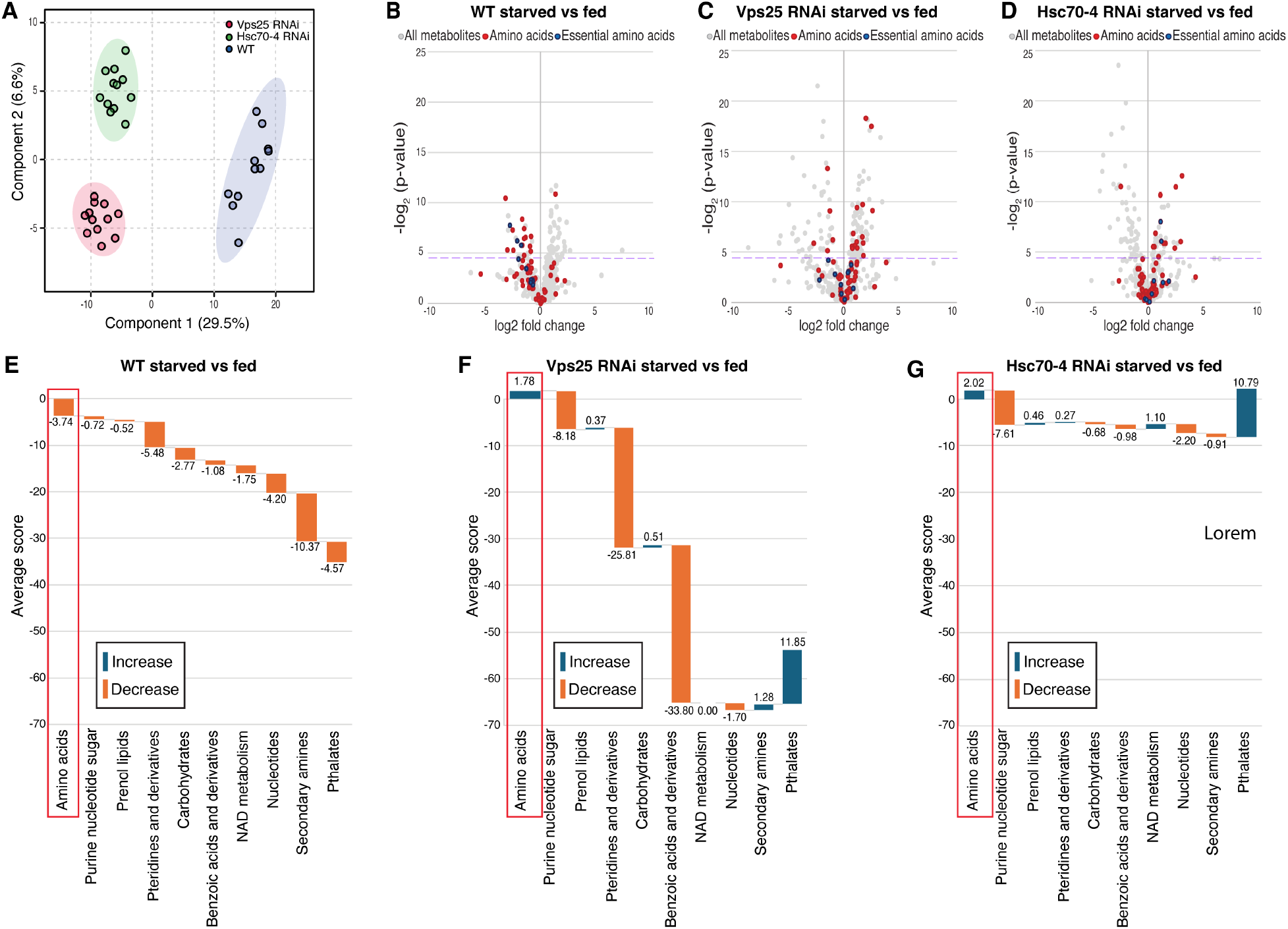
Inhibition of e-MI reverts the decrease of amino acid levels in the fat body induced by starvation. **(A)** Partial least squares discriminant analysis (PLS-DA) score plot of metabolomic profiles for WT, *Vps25* (ESCRTII) and *Hsc70-4 RNAi* shows clear separation of metabolome analyses. Each data point represents one of the six biological replicates per condition and genotype. Ellipses indicate 95 percent confidence intervals. **(B-D)** Volcano plots of differential metabolite abundance comparing FB under starved versus fed conditions for WT (B), *Vps25*RNAi (C), and *Hsc70-4 RNAi* (D), respectively. p-value cutoff >4.32 (dashed line; -log_2_-transformed). Blue dots: essential amino acids; red dots: non-essential amino acids. **(E-G)** Cumulative Waterfall plots summarizing changes in metabolite classes in the FB of starved versus fed conditions in WT (E), *Vps25* RNAi (F), and *Hsc70-4 RNAi* (G), respectively. The y-axis represents average scores (= log_2_(fold change) *-log_10_(p-value) of the metabolites classes). Positive scores indicate increased metabolite groups (blue), while negative scores indicate decreased groups (orange) within each condition.

### Integrated pathway analysis links translation downscaling to metabolic reprogramming

To understand how proteome responses correlate with the metaboIome, we performed integrated pathway analysis of the starvation-responsive metabolome and proteome using MetaboAnalyst (59). We found aminoacyl-tRNA biosynthesis and amino acid metabolic pathways, with the strongest signal observed for glycine, serine, and threonine metabolism as the most significantly affected pathways (FDR = 0.0026, impact = 1.32; Fig 7A), indicating a global perturbation in translational capacity, likely reflecting altered amino acid availability. Figure 7B shows the top 10 affected pathways grouped into four functional categories: translation-linked metabolism, amino acid metabolism, nitrogen metabolism, and carbohydrate metabolism. Again, among these, metabolism of various amino acids appeared as the dominant category, indicating extensive remodeling of amino acid pools. Aminoacyl-tRNA biosynthesis was classified as translation-linked metabolism, consistent with altered amino acid utilization affecting translational capacity. Nitrogen metabolism and glyoxylate and dicarboxylate metabolism pointed to redistribution of nitrogen and metabolic adaptation to nutrient stress. TCA cycle was included as the main carbohydrate/central carbon metabolism pathway, suggesting that carbon skeletons derived from amino acid turnover (and the provided sucrose) are funneled into mitochondrial energy production (60–62). Together, these highlighted pathways suggest that starvation-induced e-MI promotes amino acid recycling, nitrogen rebalancing, and downstream support of central carbon metabolism. Collectively, these pathway-level changes align with selective ribosomal protein degradation, suggesting that e-MI-mediated proteome downscaling supports metabolic adaptation under nutrient stress.

**Figure 7.**
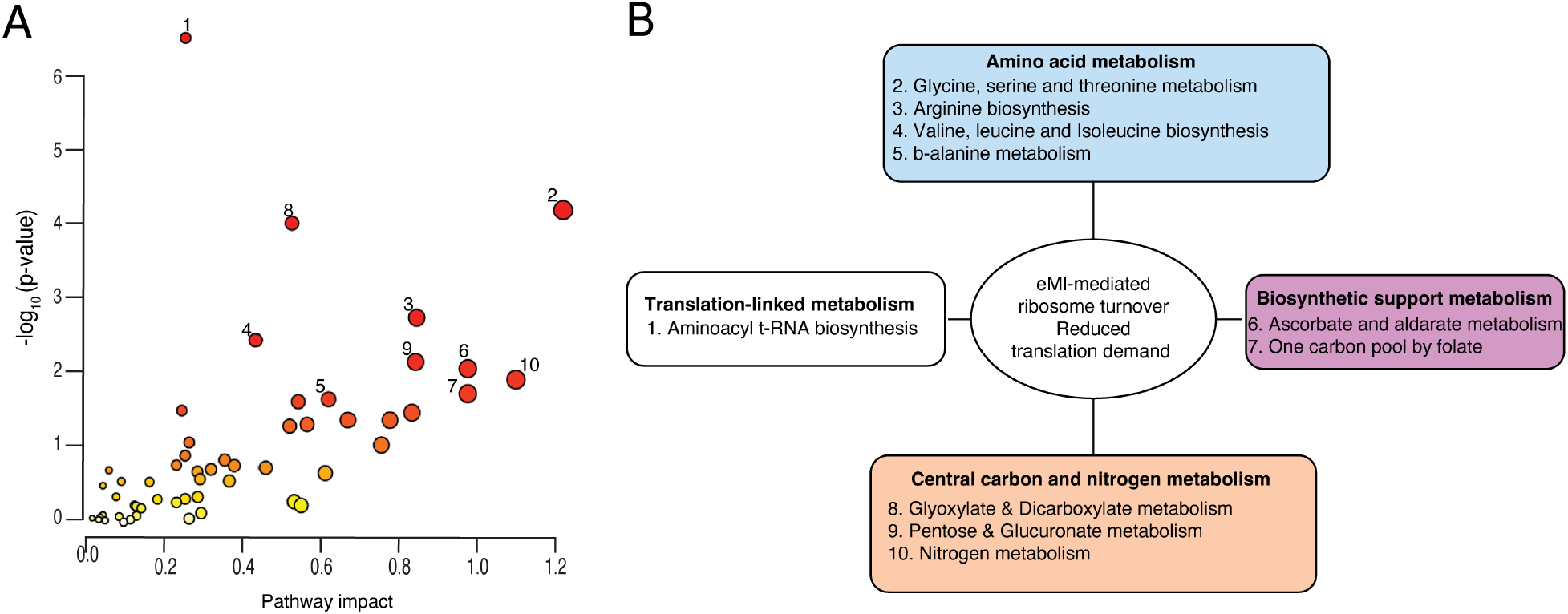
Integrated pathway analysis of proteomic and metabolomic changes in the *Drosophila* fat body in response to prolonged starvation. Joint pathway enrichment analysis using MetaboAnalyst. The 10 top hits in the dot plot analysis **(A)** were grouped according to function for the schematic in **(B)**. Each circle in (A) represents a KEGG pathway, with its impact on the x-axis and –log_10_(p value) on the y-axis. Circle size increases with pathway impact, while its color shifts from white to red with increasing -log_10_(p-value).

## DISCUSSION

Autophagy, the lysosomal degradation of cytoplasmic components is critical for many aspects of cellular homeostasis, and its loss can have detrimental consequences, particularly at old age (63). Of the three forms of autophagy, the physiological role of endosomal microautophagy is the least well understood(31). In mammals, e-MI is mostly constitutive, and HSC70/HSPA8 dependent e-MI is repressed by prolonged starvation (64, 65). Although very little is known about e-MI substrates in *Drosophila*, e-MI in flies can be constitutive such as in the larval neuromuscular junction, where it regulates the turnover of certain synaptic proteins to maintain effective synapses (25). On the other hand, in the larval FB, e-MI is induced upon exposure to prolonged times of stresses including starvation and DNA damaging agents (24, 30). The finding that starvation for times significantly longer are required for e-MI induction than what is required to elicit a maximal macroautophagic response (24, 30, 36, 37), allowed us to systematically identify likely e-MI substrates that are regulated by starvation using mass spectrometry of genetically altered FB lacking e-MI upon knockdown of *Hsc70-4* or the ESCRT III component *Vps32*. We further reasoned that (computationally) combining the analyses of proteins stabilized by *Hsc70-4* and *Vps32* knockdown upon prolonged starvation with proteins degraded at this time in WT and with prematurely degraded proteins upon overexpression of WASp (40) would allow for more robust identification of e-MI substrates.

Starving wildtype larvae by removing yeast from the food and providing only sucrose as carbon source indicated strong metabolic adaptation with translation components reduced (Fig. 1D,E) and fatty acid metabolism increased (Fig. 1F,G), possibly reflecting use of lipids stored in the FB and downregulation of TAG synthesis (66). This is consistent with studies that showed that nutrient limitation constrains energy availability, resulting in the suppression of protein synthesis, as translation is the most energy-intensive process within the cell (67–70). In addition to ribosomal components themselves, degraded proteins also include translation initiation and termination factors (eIF2α, eIF3m, eRF1) and aminoacyl-tRNA synthetases (ArgRS, SrpRα), the connection of which is also reflected as a dense network of proteins in string analyses (Fig. 1D). Also consistent with this switch upon starvation is the accumulation of a distinct protein network related to mitochondrial function such as ATP synthase subunits (ATPsynB, ATPsynF, ATPsynγ), and electron transport chain components (ND-20, UQCR-Q). Additionally, the enrichment of oxidative phosphorylation, lipid metabolism, and energy generation pathways in the significantly increased protein list suggests a metabolic shift toward catabolism and efficient ATP production (71, 72). Recent studies have highlighted the importance of ribosome turnover in maintaining cellular homeostasis during stress and have suggested that multiple proteostatic pathways cooperate to regulate ribosome abundance (73, 74).Taken together, FB cells strategically reorganize their resources to rebalance their proteome around energy efficiency, to promote cell survival during nutrient deprivation.

Our combined analysis that also removed proteins the mRNAs of which were downregulated by starvation identified 153 high-confidence e-MI candidate substrates with strong enrichment for ribosomal proteins and translation-associated processes across Gene Ontology, KEGG, and Reactome analyses (Fig. 4A–C). Lysosomal pathways were also enriched, consistent with the degradative route of e-MI (22, 24, 64, 65). In our study of e-MI substrates, translation-related processes, including initiation, co-translational targeting, and ribosome-associated quality control, were significantly enriched, suggesting that e-MI targets core components of the translational machinery. Ribosome biogenesis and protein synthesis are among the most energetically and metabolically demanding cellular processes (75, 76) and their downregulation is a hallmark of the starvation response (77, 78). This is also consistent with downregulation TOR signaling (e.g. upon starvation) leading to reduced ribosome biogenesis, translation initiation and activation of energy conservation pathways (79–82). Previous studies in mammalian cells have shown that selective autophagic pathways including e-MI can regulate translational capacity by promoting the selective degradation of ribosomal components at baseline or during nutritional stress (64, 69, 83). Consistent with this, our data support that reduction in ribosomal protein abundance leads to altered translation during nutrient stress, thereby reducing anabolic demands and reallocating resources (4, 68, 74, 84).

It has recently been shown that HSC70 dependent e-MI activity declines with age and leads to accumulation of damaged proteins (64), which highlights the importance of e-MI in maintaining proteostasis and suggest that its dysfunction contributes to age-associated proteome imbalance. However, in mammals, chaperone-mediated autophagy contributes to selective protein turnover during prolonged starvation (16), while e-MI activity is reduced during prolonged starvation (65). This divergence of e-MI activity may reflect organism-specific adaptation, particularly, given the absence of CMA in *Drosophila* (38) and confirms that e-MI can fulfill broader or functionally distinct roles depending on the autophagic landscape as was previously speculated (24). Consistently, it recently has been shown that e-MI can compensate for the lack of CMA, which leads to rerouting of a subset of CMA substrates enriched in proteins involved in translation and protein quality control to e-MI (65). A recent study of CMA substrates showed that multiple proteins of translation initiation are substrates for CMA (85). Additionally, the enrichment of ribosomal and translation-associated proteins among e-MI substrates in both, mammalian cells (65) as well as in *Drosophila* (this study) suggests a conserved role for this pathway in the selective regulation of the translation machinery to modulate protein synthesis.

Another important conclusion from our study is that e-MI-dependent proteome remodeling is genetically and functionally distinct from macroautophagy. Although starvation activates both pathways albeit with consecutive timing in *Drosophila*, we found that inhibition of MA via Atg7 depletion produced proteomic signatures enriched for amino acid degradation, fatty acid metabolism, peroxisome function, oxidative phosphorylation, and ER protein processing pathways consistent with metabolic stress and compensatory responses to impaired autophagic flux. Significantly, ribosomal protein turnover is affected by inhibition of MA as well as e-MI. Interestingly though, among the 1309 proteins called in both studies, there is apparent selectivity for the degradation of proteins by MA and e-MI, as we find evidence only for 9 proteins to be substrates for both forms of autophagy while 57 and 244 substrates seemed specific for e-MI and MA, respectively. In case of the subset of ribosomal proteins, 14 were selective e-MI and 12 were selective MA substrates, while only 2 were in common to both (Fig. 5H, I; see also Supplementary table 2). The latter is particularly surprising, as in yeast, nitrogen starvation induced ribophagy, a selective form of MA, degrades intact, whole 60S large and 40S small ribosomal subunits (68). Nevertheless, our findings are consistent with studies of e-MI and CMA in mammals that also find selective degradation of ribosomal subunits (64, 85). Future experiments will have to address how and why distinct subunits are substrates for different forms of autophagy.

Consistent with the removal of protein sources during starvation, the reduction of translation related proteins, and the enrichment of metabolic terms related to amino acid metabolism in the proteomic analysis (Fig. 1), our direct metabolomic analyses revealed significant depletion of amino acids, both including essential and non-essential ones upon prolonged starvation (Fig. 6B,E). Ribosome turnover to contributes to intracellular amino acid recycling to support metabolic homeostasis during nutrient limitation (86). The observed reduction in free amino acid levels by metabolomics suggests that autophagy-derived amino acids are rapidly utilized rather than accumulated, likely reflecting increased metabolic demand under starvation conditions. The amino acid depletion in WT FB parallels the MS data showing coordinated upregulation of mitochondrial and oxidative metabolism pathways, which likely reflects an increase in fatty acid oxidation, a well-known response to starvation in mammalian cells (87, 88). It remains to be seen why inactivation of the ESCRT machinery or Hsc70-4 partially reverts the starvation response to increase amino acid levels. Clearly, integrated pathway analysis identified aminoacyl-tRNA biosynthesis and amino acid metabolism as the most significantly enriched pathways (Fig. 7A,B) in both metabolomic and proteomic datasets together and points to a highly coordinated response to starvation involving e-MI.

The coupling between proteostasis and metabolism is increasingly recognized as a key principle of cellular stress adaptation. Autophagy-mediated protein degradation not only removes cellular components but also generates amino acids and metabolites that can be reused for energy production and biosynthetic processes (6). At the same time, selective degradation of energy-intensive cellular structures such as ribosomes can reduce biosynthetic demand. Our results suggest that e-MI contributes to this coordination by linking proteome remodeling with metabolic reprogramming during starvation. Together, our findings provide the first comprehensive *in vivo* characterization of endogenous e-MI substrates in *Drosophila* and reveal ribosome turnover as a major physiological function of this pathway. These results position e-MI as an important regulator of translational capacity and cellular homeostasis and highlight its potential role in broader stress-response pathways.

## Supporting information

Supplementary table 2 Proteome analyses

Supplementary table 3 transcriptome analyses

Supplementary table 4 Metabolome data

## Acknowledgments

We thank Drs. T.P. Neufeld (University of Minneapolis) and the Vienna *Drosophila* Resource Center for kindly sharing fly strains and Jonathan Handy for comments on the manuscript. This work was supported by AHA predoctoral fellowship # 915707 (to P.J.) and NIH/NIGMS grant GM119160 (to A.J.). The Sidoli lab is funded by the Hevolution Foundation (AFAR), the ERCM Center for AIDS Research, the NIAID (1U19AI181977), the Einstein-Mount Sinai Diabetes center, and the NIH Office of the Director (S10OD030286). The Einstein Analytical Imaging and Metabolomics Facilities are supported by NCI/P30CA013330, SIG#1S10OD023591-01 and NIH/NIDDK/ P30DK020541 and 1S10 OD021798-01A1, respectively.

## Conflict of Interest

The authors have no conflicts of interest.

## Data Availability

Proteomics raw files are available on the ProteomeXchange (PRIDE) repository under the project number PXD076298.

The metabolomics raw data are available at the NIH Common Fund’s National Metabolomics Data Repository (NMDR) website, the Metabolomics Workbench (https://www.metabolomicsworkbench.org) (89), where it has been assigned Study ID ST005003. The data can be accessed directly via its Project DOI: http://dx.doi.org/10.21228/M8H856.

## Author contribution

PJ, AJ, SiSi conceptualization; PJ, SaSu, JA, SiSi, AJ methodology and validation; PJ, JA, SiSi, formal analysis; PJ, JA investigation: PJ writing–original draft; PJ, SiSi, AJ writing–review & editing; PJ, SiSi, AJ funding acquisition.

---

**Supplementary Figure S1:**
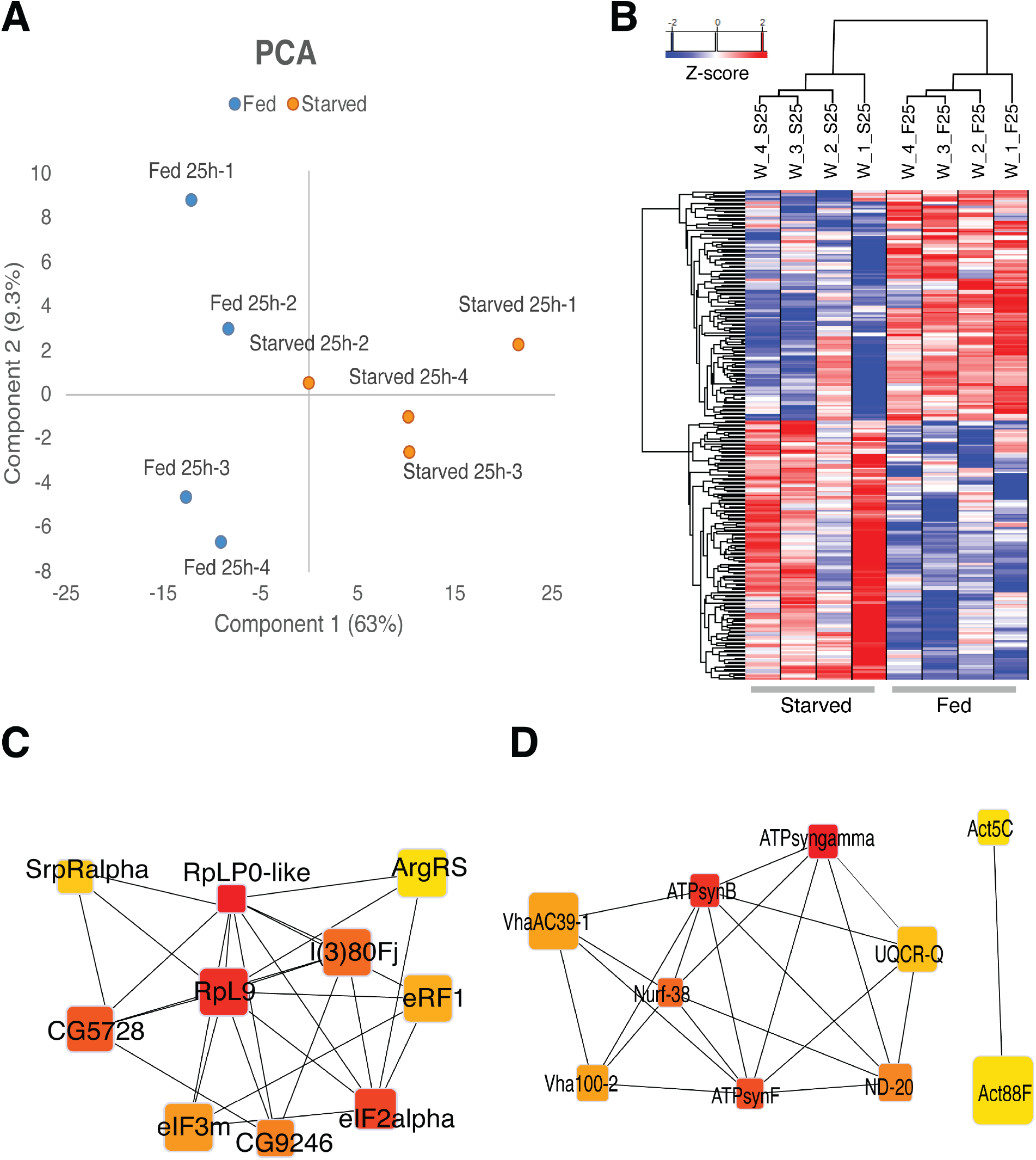
**(A)** Principal component analysis (PCA) of fed and starved wildtype larval FB samples based on normalized protein abundance profiles after 25 h of treatment. Blue: fed; orange: starved. (**B)** Heatmap clustering of all significantly changing proteins (-log_2_(p)>3) across biological replicates under fed and starved conditions showing clear segregation between starved and fed samples. Starved sample 2 was omitted from final analysis as outlier. **(C, D)** STRING network of proteins which are significantly decreased (C) or increased (D) upon starvation showing the top 10 hub proteins ranked by maximal clique centrality (MCC) using the CytoHubba plugin in Cytoscape. Each hub represents core components of ribosomal subunits, translational initiation, and aminoacyl-tRNA synthesis in (C) and subunits of ATP synthase, V-ATPase, and mitochondrial electron-transport complexes in (D). Node size reflects the fold change of each protein, and color intensity corresponds to the MCC score.

**Supplementary Figure S2:**
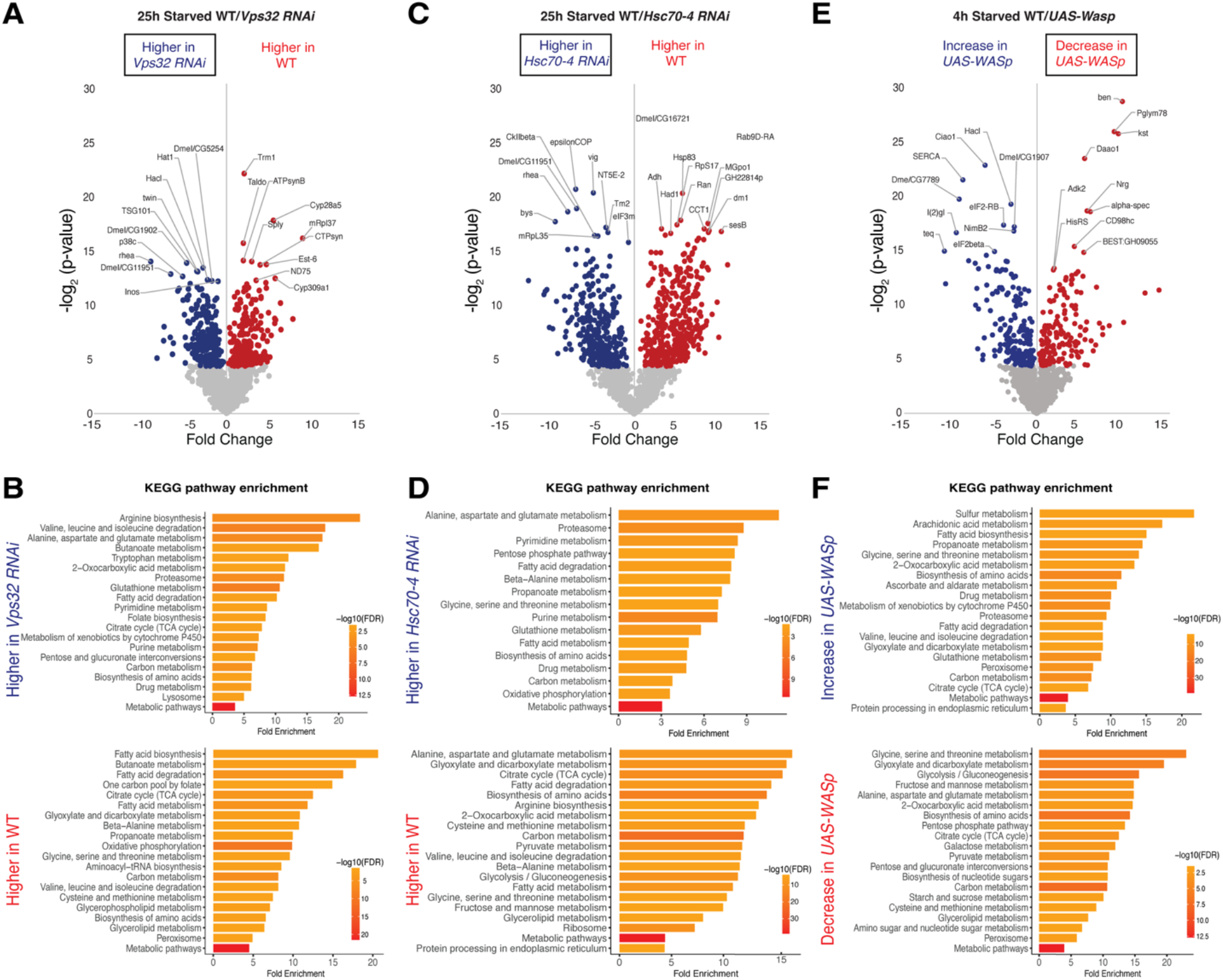
Volcano plots and KEGG pathway analyses comparing protein abundance between fed and starved larval FBs of WT with knockdown of *Vps32* **(A, B)**, *Hsc70-4 RNAi* **(C, D)**, and overexpression of WASp **(E, F)**, respectively. For the volcano plots, the x-axis shows fold change in protein abundance, and the y-axis shows statistical significance (-log_2_(p). Proteins significantly changed in abundance are colored (*p* <0.05 (>4.32 when -log_2_ transformed). Scenarios where e-MI substrates are expected are boxed. Bar plots of KEGG pathway enrichment analyses (ShinyGo) show the top 10 pathways for proteins that are significantly stabilized (upper panel) or reduced (lower panel), respectively, by knockdown or overexpression of the indicated gene. The x-axis shows ranked fold enrichment. Bar color represents statistical significance as -log_10_(FDR), with darker colors indicating higher significance (legends on the right).

**Supplementary Figure S3.**
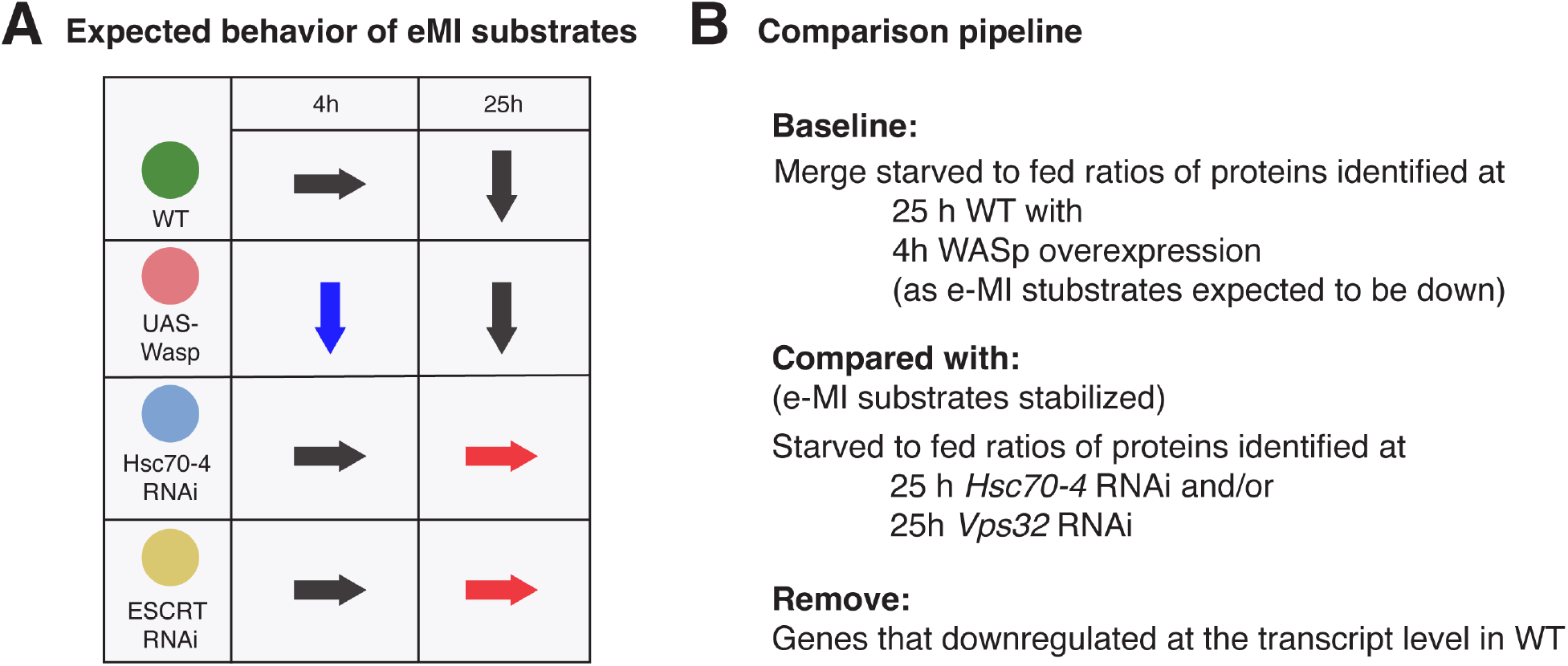
**(A)** Anticipated changes in protein levels for e-MI substrates under indicated conditions. In wildtype larva, e-MI substrates will be degraded only upon prolonged starvation (i.e. down arrow at 25h). Their degradation should be prevented by inhibition of e-MI upon RNAi mediated knockdown of *Hsc70-4* or *Vps32* (red arrows). Conversely, they should be prematurely degraded by overexpression *of WASp* at 4h of starvation (blue arrow). **(B)**. Data analysis strategy for systematic identification of potential e-MI substrates.

## Abbreviations

AGC: Automatic Gain Control
CMA: chaperone mediated autophagy
CV: coefficient of variation
DDA: data-dependent acquisition
ESCRT: endosomal sorting complex required for transport
FB: fat body
GO: gene ontology
HSC70/HSPA8: heat shock cognate protein 70
KEGG: Kyoto Encyclopedia of Genes and Genomes
LAMP/Lamp: lysosome associated protein
LE: late endosome
(e-)MI: (endosomal) microautophagy
MA: macroautophagy
MCC: maximal clique centrality
MS: Mass spectrometry
MVB: multivesicular body.
WASp: Wiscott Aldrich Syndrome protein.

## Notes

### Competing Interest Statement

The authors have declared no competing interest.

