## Supplementary material for "Ribosomal proteins are major substrates of starvation-induced endosomal microautophagy in *Drosophila*": Suppl Table 1

**Supplementary Table 1:** Internal standards in the extraction solution for metabolomics

| Name | Concentration | Name | Concentration |
| --- | --- | --- | --- |
| ^13^C_4_-Succinic acid | 0.18 ug/mL | ^13^C_6_-Citric acid | 0.6 ug/mL |
| ^15^N_5_-AMP | 1.06 ug/mL | d_5_- Indoleacetic acid | 0.27 ug/mL |
| d_5_- Tryptophan | 0.3 ug/mL | d_3_- L-Carnitine | 0.06 ug/mL |
| d_3_-Acetyl- L-carnitine | 0.06 ug/mL | d_3_-Butyryl-L-carnitine | 0.06 ug/mL |
| d_9_-Isovaleryl-DL-carnitine | 0.06 ug/mL | d_3_-Hexanoyl-L-carnitine | 0.06 ug/mL |
| d_3_-Octanoyl-L-carnitine | 0.06 ug/mL | d_3_-Decanoyl-L-carnitine | 0.06 ug/mL |
| d_3_-Hexadecanoyl-L-carnitine | 0.06 ug/mL | d_3_-Octadecanoyl-L-carnitine | 0.06 ug/mL |
